# A *de novo* approach to disentangle partner identity and function in holobiont systems

**DOI:** 10.1101/221424

**Authors:** Arnaud Meng, Camille Marchet, Erwan Corre, Pierre Peterlongo, Adriana Alberti, Corinne Da Silva, Patrick Wincker, Eric Pelletier, Ian Probert, Johan Decelle, Stéphane Le Crom, Fabrice Not, Lucie Bittner

## Abstract

**Background:** Study of meta-transcriptomic datasets involving non-model organisms represents bioinformatic challenges. The production of chimeric sequences and our inability to distinguish the taxonomic origins of the sequences produced are inherent and recurrent difficulties in *de novo* assembly analyses. The study of holobiont transcriptomes shares similarities with meta-transcriptomic, and hence, is also affected by challenges invoked above. Here we propose an innovative approach to tackle such difficulties which was applied to the study of marine holobiont models as a proof of concept.

**Results:** We considered three holobionts models, of which two transcriptomes were previously assembled and published, and a yet unpublished transcriptome, to analyze their raw reads and assign them to the host and/or to the symbiont(s) using Short Read Connector, a k-mer based similarity method. We were able to define four distinct categories of reads for each holobiont transcriptome: host reads, symbiont reads, shared reads and unassigned reads. The result of the independent assemblies for each category within a transcriptome led to a significant diminution of *de novo* assembled chimeras compared to classical assembly methods. Combining independent functional and taxonomic annotations of each partner’s transcriptome is particularly convenient to explore the functional diversity of an holobiont. Finally, our strategy allowed to propose new functional annotations for two well-studied holobionts and a first transcriptome from a planktonic Radiolaria-Dinophyta system forming widespread symbiotic association for which our knowledge is limited. Conclusions

In contrast to classical assembly approaches, our bioinformatic strategy not only allows biologists to studying separately host and symbiont data from a holobiont mixture, but also generates improved transcriptome assemblies. The use of Short Read Connector has proven to be an effective way to tackle meta-transcriptomic challenges to study holobiont systems composed of either well-studied or poorly characterized symbiotic lineages such as the newly sequenced marine plankton Radiolaria-Dinophyta symbiosis and ultimately expand our knowledge about these marine symbiotic associations.

## Background

In its scientific acceptation, symbiosis is defined as the living together of unlike organisms whatever the nature of their relationship [1], ranging from parasitism to mutualism. Symbiosis is a widespread phenomenon in the biosphere and plays crucial roles in evolution and ecology. One of the most popular examples of mutualism is the interaction between fungi and land plants, where fungi form mycorrhizae that help land plants to retrieve nutrients from soil [2]. In the ocean, benthic coastal ecosystems are structured and supported by symbiotic associations involving multipartners such as corals (Cnidaria, i.e. multicellular eukaryotes), microalgae (Dinophyceae, *Symbiodinium* spp., *i.e*. unicellular eukaryotes), and Bacteria. Breakdown of this symbiosis ultimately leads to coral bleaching (the loss of photosynthetic symbionts), dramatically affecting the whole reef ecosystems [3]. While coral bleaching has been largely studied, there is a growing evidence that other partners are involved in the holobiont system, and contribute to make coral reef persisting in oligotrophic seas. For instance, symbiotic association between sponges (Porifera, i.e. multicellular eukaryotes) and Bacteria (prokaryotes) allows Bacteria to grow within the mesohyl matrix of the sponge where they can be metabolically active and persist in a highly oligotrophic habitat. The symbiotic interactions between sponges and bacteria are currently poorly understood from the genomic point of view [4]. Symbiotic associations involving two unicellular eukaryotes are also widespread in the oceanic plankton [5–7,9]. For instance, the cosmopolitan mutualistic associations between heterotroph Radiolaria (host) and endosymbiotic microalgae play significant ecological and biogeochemical roles in the oceans [8] but the underlying genomic basis of such associations remains uncharacterized. Although not cultivable *in vitro*, nucleic acids extraction is nevertheless possible on such symbiotic partnerships, and this recently allowed shedding light on the identity of the partners and their co-evolutionary history [6, 7]. Several symbiotic microalgae have been identified using such molecular approaches, and many of them belong to the eukaryote Dinophyta [9]. Mainly because of their highly complex and large genomes, the lack of reference genomes for both Dinophyta and Radiolaria make their study challenging for *de novo* assembly and functional annotation [10, 11]. The study of the RNA mixture from a holobiont system, being composed of the host and its symbiotic microbial communities offers the opportunity to characterized functional aspects through their expressed genes, and so in different abiotic conditions/decoupling the functional/metabolic role of each partner.

Currently, RNA-seq approaches are the best available tools to obtain large amount of genomic information from uncultured organisms isolated in the environment [12, 13]. RNA sequencing for a holobiont is now possible [14–16] and has promoted the development of sequencing projects [17] for non-model organisms. Non-model holobiont RNA-seq datasets corresponds to a mixture of data coming simultaneously from the host and from the symbiont(s). Studying such datasets share similarities with meta-transcriptomics and requires *de novo* assembly of transcripts sequences, which implies large computational resources and has the potential to introduce biases such as generating numerous chimeric sequences resulting from the mis-assembly of RNA fragments from the host and from the symbiont(s) [18, 19]. A variety of analysis strategies has been developed to address meta-transcriptomic challenges. Some of these strategies avoid the assembly step to focus on identifying abundant species and significant functional differences between meta-transcriptomes directly from raw data [20, 21]. Other strategies use statistical tools and machine learning algorithms to improve the quality of *de novo* assembly of meta-transcriptome by learning from their abundance information [22].

Here we developed an original strategy aiming at improving *de novo* assembly for newly generated holobiont sequence dataset. We chose to use the Short Read Connector software in its Counter version (SRC_c) [23]. SRC_c is a fast kmer-based method initially developed to estimate the similarity between numerous (meta)genomic datasets by extracting their common sequences. We focused on holobiont transcriptomes for which *a priori* no or little genomic knowledge has been previously produced for host and symbionts, and we used SRC_c to compare these holobiont sequences to publicly available databases. Our strategy is to use SRC_c to assign at best holobiont sequences either to the host or to the symbionts before the *de novo* assembly step (Fig. 1). It allows then independent assembly of the datasets and prevents the potential mis-assemblies of reads from diverse origin.

**Figure 1.**
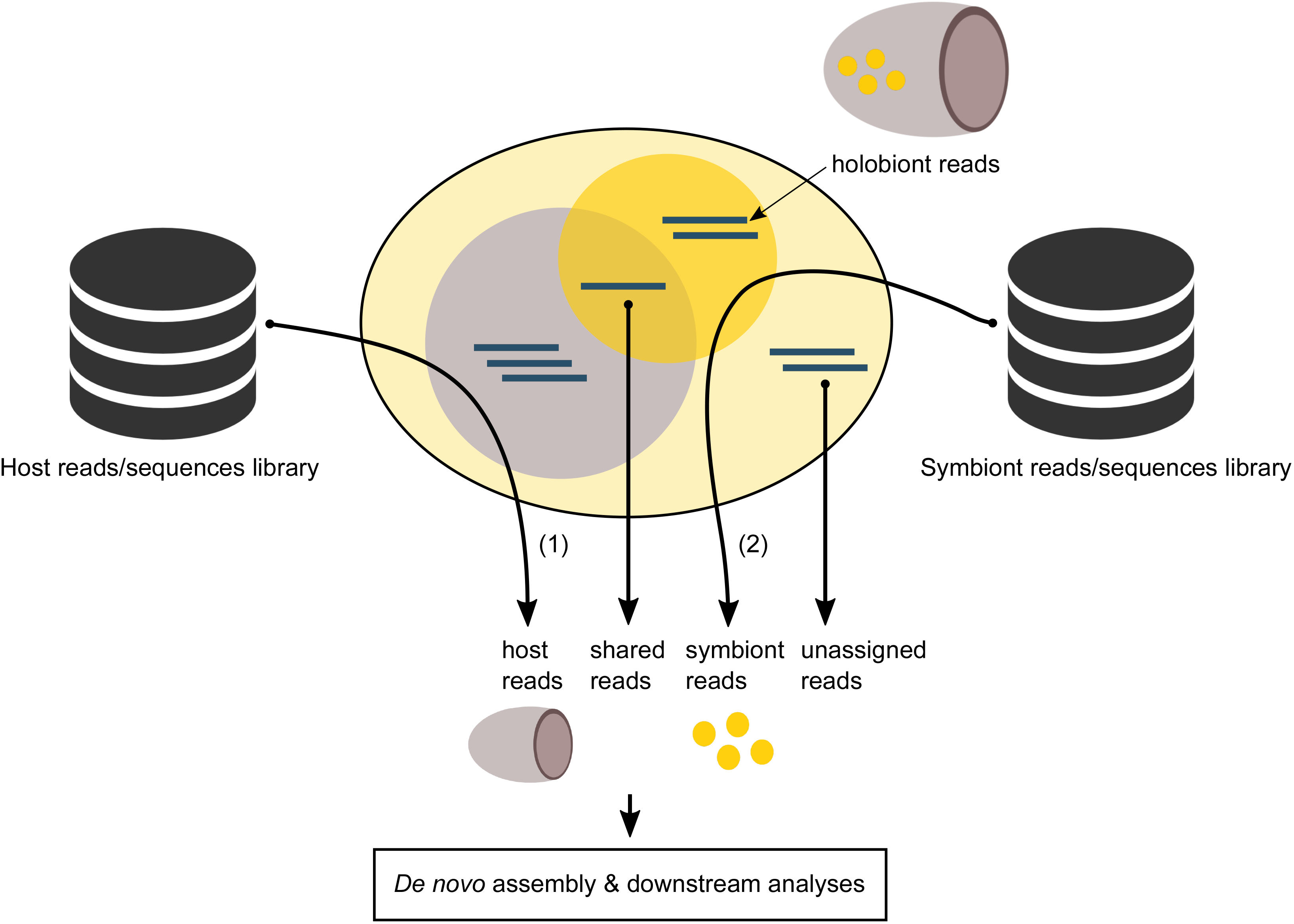
Theoretical overview on the application of SRC_c on holobiont transcriptome. The comparisons to (1) host and (2) symbiont reads/sequences library are done against the entire holobiont dataset to retrieve host and symbiont similar reads. The 4 resulting subsets (host, symbiont, shared and unassigned reads) are then processed independently (de novo assembly and downstream analyses)

We applied our strategy to disentangle the sequences and then *de novo* assemble the transcriptome of three distinct marine holobiont systems (Fig 2). Two of them were already assembled and published. The first model (M1) involves a Cnidaria host (*Orbicella faveolata*, belonging to the Metazoa) and Dinophyta symbionts (*Symbiodinium* spp., a unicellular eukaryote belonging to the Alveolata) forming a mutualistic association [24, 25]. This symbiotic association represents the best-known example of symbiosis in marine ecosystems, and many studies have been made trying to understand coral bleaching events (*i.e*. the loss of symbionts) [26, 27]. The coral holobiont also encompass other microorganisms consisting of bacteria, archaea, fungi, viruses [28, 29]. In the second holobiont model (M2) the marine sponge *Xestospongia muta* (Porifera) harbors a dense (40% of its volume) and diverse microbial community including marine protists (*e.g*. fungi), archaea and mainly bacteria [30–32]. The symbiotic associations between sponges and bacteria (suggested to be commensalism [33]) have become a major research focus to understand how sponges and their microbial communities can perform a variety of functional roles such as nutrition, cycling of metabolites and host defense allowing them to proliferate in oligotrophic conditions [34, 35]. We chose a third, yet unpublished, holobiont dataset (M3) involving two distinct lineages of protists (unicellular eukaryotes): the radiolarian *Collozoum* sp. as host and Dinophyta symbionts belonging to the *Brandtodinium nutricula* species [6]). In this association, the radiolarian host forms a gelatinous matrix of several centimeters, which contains hundreds of host cells and thousands of symbiotic microalgae (refer to image). Recent studies showed that this symbiosis is widely distributed in the ocean and significantly contribute to biomass and carbon export in the open ocean [36, 37]. As a proof of concept, we these holobiont transcriptomes datasets, and we compared quantitatively and qualitatively results obtained when involving SRC_c or not.

**Figure 2.**
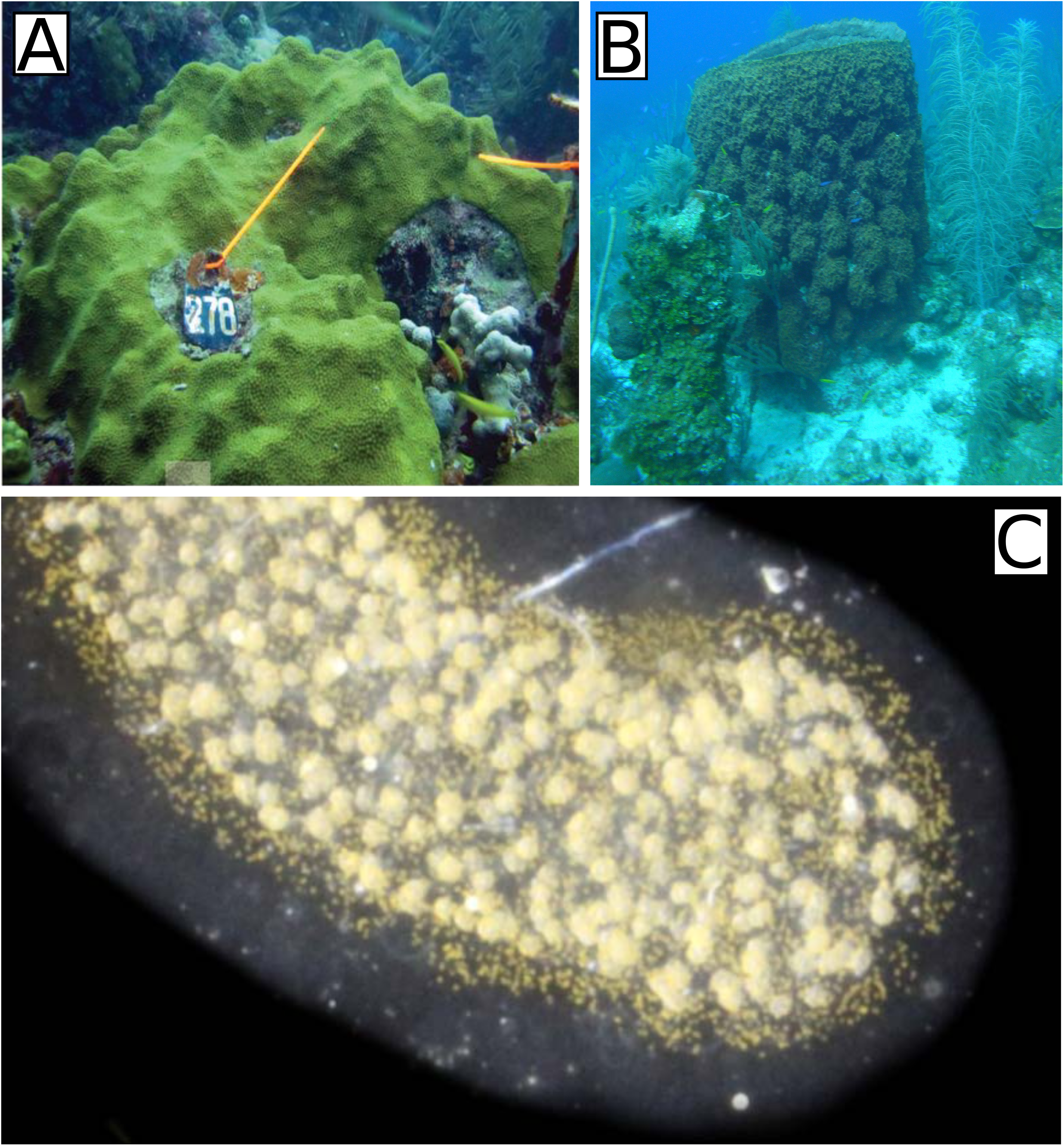
Pictures of the 3 holobiont models. (A) the *Orbicella faveolata* holobiont in symbiosis (unbleached) in 2010 at reefs of La Parguera, Puerto Rico (credits: [24]). (B) A *Xestospongia muta* specimen in symbiosis on a coral reef near Little Cayman in the Caribbean (credits: Cara Fiore, january 14, 2015 http://feedthedatamonster.com). (C) A Collodaria colony with symbionts sampled in South Pacific Ocean at station 112.01 of the Tara Pacific expedition in 2011 (credits: Johan Decelle).

## Results

### Choice of holobiont models and building of host and symbiont reference libraries

For each of the three holobiont models (Fig. 2), we built reference sequences libraries representing host and symbiont(s) by selecting the taxonomically closest organisms available in public datasets (see Methods, Additional files 1). The M1 host reference library encompasses 22 assembled transcriptomes from Cnidaria (including data from the host species *Orbicella faveolata* itself) and the M1 symbiont reference library encompasses 123 RNA-seq reads datasets (including the presumed major symbiont *Symbiodinium* spp. [38]). The M2 host reference library involves 4 RNA-seq reads datasets from distinct Porifera genera (and differ from the *Xestospongia* genus) whereas the M2 symbiont reference library corresponds to the *Tara* Oceans metagenomic gene catalogue (OM-RGC) assembled from the pico-planktonic fractions (< 3 μm) including bacteria or Archaea [39]. For M3, we used the four Rhizaria transcriptomes published so far to create the reference host library whereas the same library as for M2 has been used for symbiont references. All reference libraries described above include assembled transcriptomes, genomes or RNA-seq raw reads datasets for eukaryotic or prokaryotic holobiont partners (Additional files 1). Their sizes vary from 4.5 Mbp to 25 Gbp with sequences length from 100 bp to 84 Kbp (Additional files 1).

### Disentangling the holobiont sequences

Disentangling the holobiont sequences for all three models (M1, M2 and M3), the SRC_c memory footprint was far lower than our cluster’s capacity (Tab. 1), even for the biggest data set to index (M2 symbiont library of 25 Gbp has been built with 58.9G of RAM). This induces that any addition of data can be considered.

**Table 1.** Performances of SRC_c

|  |  | Time(hh:mm:ss) | Memory (Gb) |
| --- | --- | --- | --- |
| **Cnidaria-Dinophyta holobiont (M1)** | all symbionts library (M1a) | 15:40:42 | 34,2 |
|  | *Symbiodinium* spp. library (M1b) | 01:34:57 | 6,96 |
|  | other symbionts library (M1c) | 15:08:45 | 33,7 |
|  | host library | 01:06:56 | 3,9 |
| **Porifera-Bacteria holobiont (M2)** | symbionts library | 21:04:47 | 58,9 |
|  | host library | 02:46:06 | 9,60 |
| **Radiolaria-Dinophyta holobiont (M3)** | symbionts library | 07:05:28 | 4,10 |
|  | host library | 00:05:57 | 3,9 |

Memory peak and wallclock time of SRC\_c indexing and query steps on the several data sets for models M1, M2 and M3.

The comparison of holobiont reads to reference host and symbiont sequence libraries enabled to identify and classify them into four categories (Fig. 1): (1) reads specific to the host, (2) reads specific to the symbionts (including microalgae, bacteria…), (3) reads which can be assigned to both reference libraries and (4) reads which do not match any reference library (referred as to ‘unassigned’). For the three holobiont models, the distribution within the four categories is reported in Tab. 2.

**Table 2.** SRC_c assignment results for the holobiont models M1, M2 and M3

|  |  | # reads | % reads from holobiont |
| --- | --- | --- | --- |
| *Orbicella faveolata* holobiont (M1a) | **total** | **775 025 024** |  |
|  | assigned to host library | 498 008 661 | 64.26% |
|  | assigned to symbiont library | 56 011 798 | 7.23% |
|  | shared | 32 133 818 | 4.15% |
|  | unassigned | 188 870 747 | 24.37% |
| *Orbicella faveolata* holobiont (M1b) | assigned to host library | 500 145 229 | 64.53% |
|  | assigned to symbiont library | 54 850 148 | 7.08% |
|  | shared | 29 997 250 | 3.87% |
|  | unassigned | 190 032 397 | 24.52% |
| *Orbicella faveolata* holobiont (M1c) | assigned to host library | 521 591 231 | 67.30% |
|  | assigned to symbiont library | 4 817 450 | 0.62% |
|  | shared | 8 551 248 | 1.10% |
|  | unassigned | 240 065 095 | 30.98% |
| *Xestospongia muta* holobiont (M2) | **total** | **33 220 038** |  |
|  | assigned to host library | 6 193 678 | 19.04% |
|  | assigned to symbiont library | 825 154 | 10.64% |
|  | shared | 5 112 031 | 8.63% |
|  | unassigned | 21 090 174 | 61.69% |
| *Collozoum* sp. holobiont (M3) | **total** | **97 957 794** |  |
|  | assigned to host library | 3 188 944 | 3.26% |
|  | assigned to symbiont library | 23 234 402 | 23.72% |
|  | shared | 531 432 | 0.54% |
|  | unassigned | 71 003 016 | 72.48% |

SRC\_c assignment results for the Cnidaria-Dinophyta holobiont model (M1) against the complete Dinophyta library (M1a), the *Symbiodinium* spp. exclusive library (M1b) and the Dinophyta library excluding *Symbiodinium* spp. (M1c), the Porifera-Bacteria holobiont model (M2) and the Radiolaria-Dinophyta holobiont model (M3).

With M1, SRC_c assigned 64.3% of the holobiont reads to the cnidarian host and 7.2% to the Dinophyta symbiont full library (analysis M1a, Tab. 2). Restricting the symbiont library to the genus *Symbiodinium* spp. sequences allowed obtaining similar results with 64.5% of the reads identified as specific to the host library and 7.1% as specific to the symbiont library (analysis M1b, Tab. 2). On the contrary, when *Symbiodinium* spp. is removed from the library, only 0.6% of the holobiont reads could be assigned to the symbionts and the proportion of reads assigned to the host increases up to 67.3% (analysis M1c, Tab. 2). Our tests on the symbionts library showed that the library content impacted drastically the reads retrieval by SRC_c and demonstrated the sensitivity of the strategy. Considering these results, we focused on the M1a dataset for downstream analyses. We also noticed that shared reads (i.e. found in both host and symbiont libraries) always represent the lowest proportion of holobiont reads (M1a, M2 and M3).

### De novo assembly, contigs evaluation and downstream analyses for M1 and M2

For each holobiont transcriptome, four subsets of reads were independently *de novo* assembled, producing contigs from which protein domains were then predicted and functionally annotated (Fig. 1). For holobiont models M1a and M2, the assembly metrics, statistics and functional annotations from our contigs are summarized in Tab. 3, and comparison with previous studies are shown in Fig. 3.

**Figure 3.**
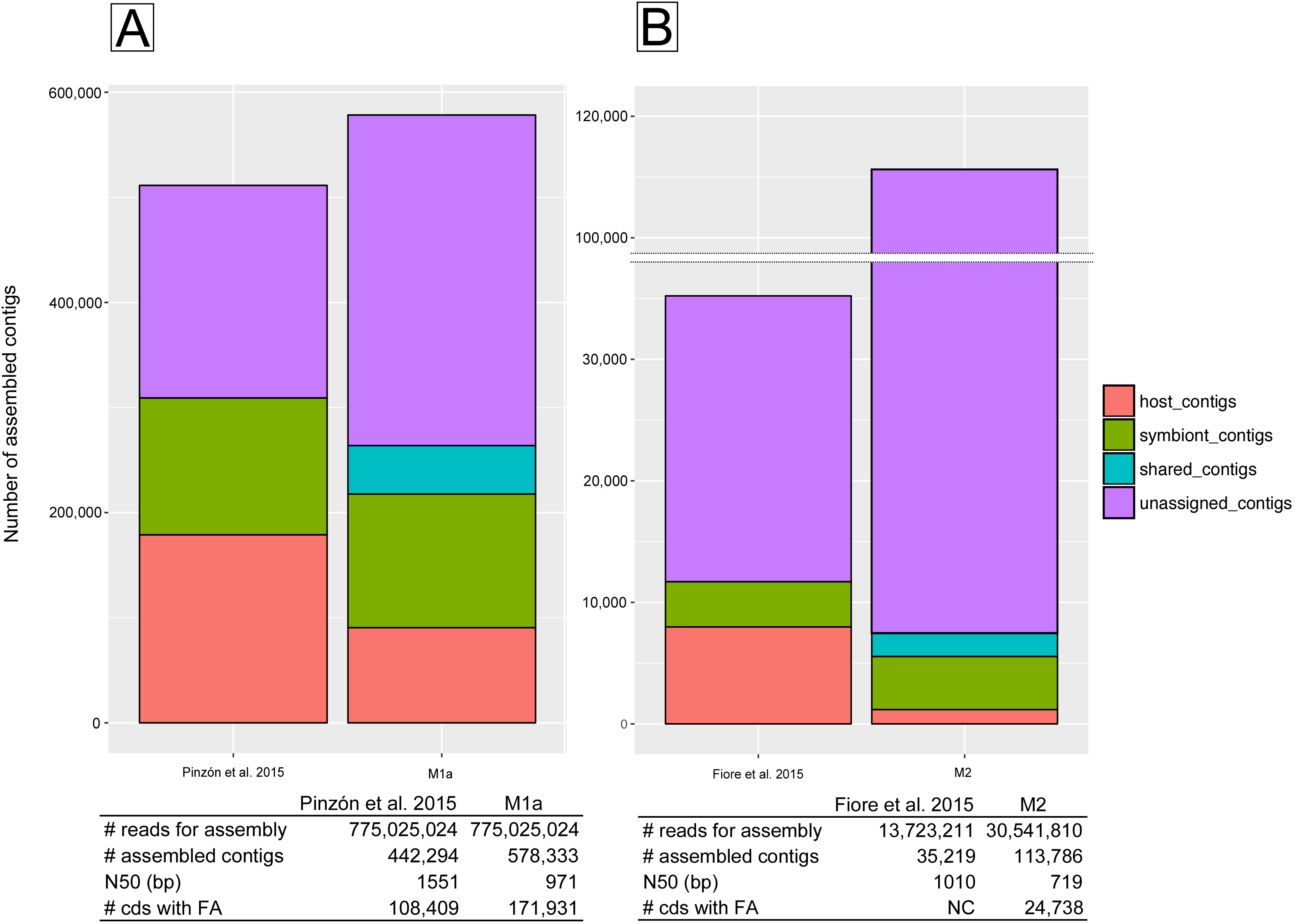
Overview and comparison to previous studies. The total assembled contigs for holobiont model M1a and M2 compared to the assembled meta-transcriptomes from (A) Pinzon et al. 2015 [24] and (B) Fiore et al. 2015 [30] respectively are shown. General details about *de novo* assembly and functional annotation (termed FA) features are presented in corresponding tables for (A) holobiont model M1a versus Pinzon et al. 2015 [24] meta-transcriptome, and (B) holobiont model M2 versus Fiore et al. 2015 [30]. NC means that exact number is not communicated.

**Table 3.**
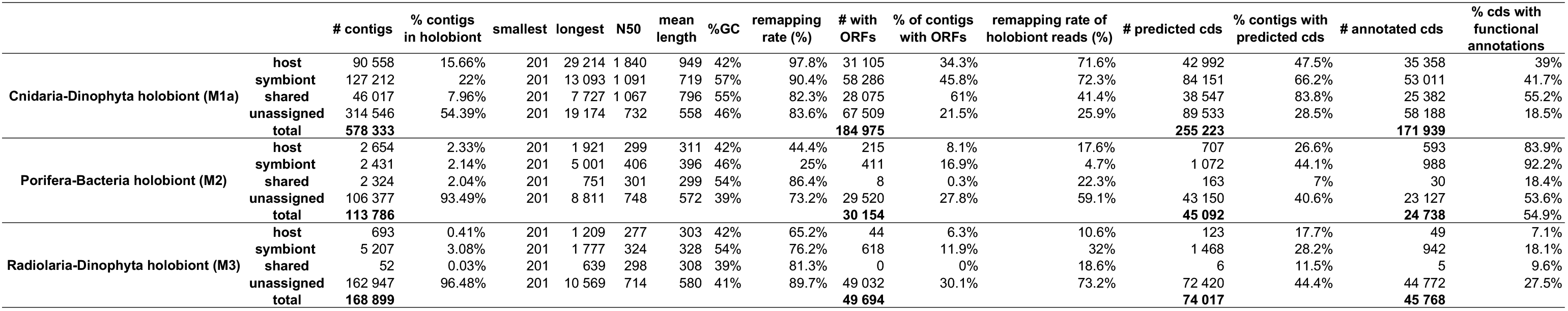
*De novo assembly* metrics and downstream analysis of SRC_c resulting subsets for holobiont models M1a, M2 and M3. (upload as additional files)

Compared to the studies where these datasets were initially published, our strategy allows considering more reads (16,818,599 reads for M2) in the assembly step as well as obtaining more assembled contigs (136,039 contigs for M1a and 78,567 contigs for M2) (Fig. 3). The contigs metrics show shorter lengths of N50 (580 bp shorter for M1a and 219 bp shorter for M2) (Fig. 3) compared to the original publication analyses. The M1a contigs display high remapping rates (>80%) while M2 contigs show mixed results (25% < x < 86%) (Tab. 3). With M1a, a total of 255,223 protein coding domains were predicted for 44.1% of the assembled contigs and functional annotations were found for nearly 30% of these protein coding domains (Tab. 3). With M2, protein coding domains were predicted for 39.6% of the contigs, and 54.9% of the domains were functionally annotated (Tab. 3). In comparison with statistics available in previous studies, we obtained 1.6 times more functionally annotated contigs for M1a (Fig. 3). This comparison for M2 could not be made since the exact number of annotated contigs in the holobiont assembly has not been reported by the authors.

To further test the usefulness of the reads sorting before the *de novo* assembly step, we compared the contigs assignment of M1a and M2 (column 1 in Tab. 3) with a taxonomic assignment performed with MEGAN6 [40]. For M1a, MEGAN6 assigned 71,143 contigs to the host *Orbicella faveolata* and 148,409 contigs to the symbiont *Symbiodinium* spp. (Additional files 2). All the contigs assigned to *Orbicella faveolata* with MEGAN6 were also found with the SRC_c strategy (Tab. 3) but we assigned 19,415 more contigs to the host category. On the contrary, MEGAN6 assigned 21,197 additional contigs to *Symbiodinium* spp. compared to our categorization strategy (Tab. 3, Additional files 2). With M2, MEGAN6 assigned 11 contigs to the host *Xestospongia muta* (Additional files 2) which is far less than the 2,654 contigs defined with the SRC_c strategy (Tab. 3). However, MEGAN6 assigned also 33,810 contigs to *Amphimedon queenslandica*, a distinct sponge species which is not supposed to be the host in this holobiont system. MEGAN6 also succeeded to assign more contigs to Bacteria (21,318 contigs) than the SRC_c strategy (2,431 contigs) (Tab. 3).

Our functional annotations were compared to initial studies having generated these datasets. As previous publications do not provide exhaustive lists of the functional annotations and their corresponding abundance, these comparisons are essentially qualitative. For the *O. faveolata* host (M1), we only found similarities in the most abundant annotations (Additional file 3). At biological processes level, both our study and Pinzón et al. 2015 found abundant metabolic process GO term (GO:0008152; 819 CDs (coding sequences) and 5,278 genes respectively). At the molecular function level, our host contigs mainly corresponded to binding protein (GO:0005515; 36,349 CDs) while Pinzón et al. 2015 mainly found catalytic activity functions (GO:0003824; 3,361 genes). For M2, rare overlaps are found between Fiore et al. 2015 and our annotations (Additional file 3): at the biological processes level, 1 of the top 15 host annotations is identical (signal transduction (GO:0007165)) and 3 of the top 15 symbiont annotations are in common (metabolic process (GO:0008152); proton transport (GO:0015992) and protein folding (GO:0006457)).

### Benchmark comparisons on M3: what difference does it make to use SRC_c?

For the holobiont model M3, assembly metrics, abundance of chimera and functional contents were compared between the SRC_c contig sets (host, symbiont, shared and unassigned) and a direct *de novo* assembled transcriptome obtained from holobiont reads considered all together (this strategy is hereafter called *noSRC*).

The assembly metrics appear very similar between SRC and noSCR (Tab. 4). A comparable number of reads were used for the assembly step and a comparable number of assembled contigs were obtained. The N50 value for the *noSRC* strategy is slightly longer while the remapping rates are 5% better with the SRC strategy. Calculation times performed on the same bioinformatic cluster revealed that the SRC strategy was 40 hours longer. The SRC strategy showed 50% less chimeras (418 contigs) than the *noSRC* strategy (777 contigs) with most chimeras contained in the unassigned set (Tab. 4). We noticed slightly less annotated CDs with the SRC strategy (45,768 against 47,260), however the number and the composition in GO annotations were very similar (Fig 1 from Additional files 4). We found 253 different biological processes with SRC against 255 with the *noSRC* strategy, and the top 5 functional annotations in the 3 Gene Ontology levels (Molecular Function, Biological Process and Cellular Component) are strictly identical (Fig 2 from Additional files 4). Considering all GO annotations, 686 are common to both strategies while 52 are exclusive to the SRC strategy and 42 to the *noSRC* strategy (Fig 3 from Additional files 4).

**Table 4.** SRC_c impact on Radiolaria-Dinophyta holobiont model (M3)

|  |  | no SRC | SRC |
| --- | --- | --- | --- |
| **# reads used in assembly** |  | 48 733 956 | 48 660 697 |
| **# assembled contigs** |  | 167 023 | 168 899 |
| **# predicted cds** |  | 75 450 | 74 017 |
| **# annotated cds** |  | 47 260 | 45 768 |
| **N50 (bp)** | **total** | 818 | 702 |
|  | host |  | 277 |
|  | symbiont |  | 324 |
|  | shared |  | 298 |
|  | unassigned |  | 714 |
| **remapping rates (%)** | **total** | 85,6 | 90,5 |
|  | host |  | 65,2 |
|  | symbiont |  | 76,2 |
|  | shared |  | 81,3 |
|  | unassigned |  | 89,7 |
| **# chimera** | **total** | 777 | 418 |
|  | host |  | 4 |
|  | symbiont |  | 47 |
|  | shared |  | 0 |
|  | unassigned |  | 367 |
| **Calculation time (min)** | **total** | 330 | 2 783 |
|  | SRC |  | 2 460 |
|  | assembly | 330 | 323 |

SRC\_c impact on assembled contigs quality and calculation times of Radiolaria-Dinophyta holobiont model (M3) compared to a direct meta-transcriptome assembly strategy. In grey are displayed the details for SRC\_c holobiont categories (host, symbiont, shared and unassigned). The “total” values for N50 and remapping rates of the SRC\_c strategy were re-calculated on pooled contigs from host, symbiont, shared and unassigned subsets.

To test the usefulness of the categorization step, all M3 contigs from the SRC strategy were taxonomically assigned using MEGAN6 (Additional files 5). MEGAN6 assigned 10 contigs to Collodaria whereas the SRC strategy assigned 683 contigs to the host category. MEGAN6 assigned 1,383 contigs to Dinophyceae compared to the 5,207 contigs categorized as symbionts. The leftover MEGAN6 contigs were assigned to Bacteria and Archeae (3,799 contigs), Viruses (76 contigs), other-eukaryotes (29,524 contigs) and 127,447 contigs remained unassigned (162,947 unassigned contigs with the categorization strategy).

## Discussion

### The use of SRC_c to tackle meta-transcriptomic challenges

The strategy proposed here is a practical and scalable solution for transcriptomic assembly of non-model holobiont organisms, from which no or limited genomic information is available. The present implementation of SRC_c [23] based on reference databases of putative partners involved in the holobiont consortium, and our analysis strategy, enabled the categorization of holobiont reads into 4 subsets. Then, these subsets have been independently assembled, limiting potential creation of chimeras while generating more assembled contigs (Fig. 1). The newly defined shared reads category represents an added value compared to other holobiont transcriptomic studies and has been later processed with the same methodology than other categories (Fig. 1).

With respect to the reference libraries, as exemplified in M1, when the expected symbiotic partner (i.e. *Symbiodinium* spp.) is missing from the reference library, the number of reads assigned to the symbiont category decreases drastically from 50M reads to nearly 5M reads (Tab. 2). The M2 and M3 libraries do not contain reference data for the expected host partner, and consequently only a low proportion of the holobiont reads are assigned to the host (19% and 3%, respectively). Accordingly, the proportion of unassigned reads is directly linked to both host and symbiont libraries content with respect to the studied holobiont. Overall, less unassigned reads were observed when the “correct” actors are involved (M1a: 24.4%) compared to the poorly studied models (M2: 61.6% and M3: 72.5%). These results highlight the sensitivity and specificity of the SRC_c requests that relies on the completeness of the database to accurately sort the reads of the holobiont. The SRC_c assignation step could be further improved by adding more sequences (*i.e*. reads, assembled genes or transcripts) from taxonomically close species to the host and symbiont reference libraries, but also from parasites and viruses that are common in multicellular and unicellular host cells.

We also compared the metrics of our SRC_c contigs to those from previous studies (M1a and M2) [24, 30]. With the SRC_c strategy, the amount of reads used for *de novo* assembly of M2 was higher than for previous studies (Fig. 3). We found that, not only our strategy allowed defining a new category of contigs (the “shared” contigs), but also allowed assembling more contigs than previous studies (Fig. 3). Our contigs metrics showed lower N50 for both models compared to previous studies, but showed higher remapping rates overall for M1a (up to 90%, (Tab. 3)). Differences in the number of contigs as well as contigs metrics could be the results of the use of distinct *de novo* assembly software: *e.g*. M2 data were processed with the CLC workbench [CLC bio, Boston, MA, USA; (https://www.qiagenbioinformatics.com/)] in the original publication while we choose the Trinity software [41] otherwise we suggest that SRC_c do not significantly impact transcriptome assembly. In fact, previous studies had shown that Trinity is able to generate more assembled contigs than the CLC assembler when applied on the same dataset. It is also known that assembled contigs from Trinity are shorter than those assembled by CLC but provided similar proportion of significant hits to the nr database [42].

With M1a, our strategy produced 1.5 times more CDs with a functional annotation (Fig. 3). At that point we are unable to tell whether this observation can be the consequence of a better suited assembly strategy (SRC_c treatment and / or assembly software), and / or the use of a different annotation pipeline, and / or the supplementation of reference annotation databases between 2015 [24] and 2017.

With M3 analyses we can estimate how SRC_c impacts the *de novo* assembly step and downstream analyses compared to a more conventional protocol (here called the *noSRC* strategy) (Tab. 4). The calculation time for the two protocols showed that the SRC_c strategy increases the total time with nearly 40 additional hours compared to a classic assembly strategy (Tab. 4). However, compared to classic strategies, the SRC_c strategy has the tremendous benefit to create directly 4 independent subsets (two of which are directly assigned to holobionts partners). Otherwise, minimal differences were found between the two protocols concerning the number of assembled contigs and, as for M1a and M2, the SRC_c strategy produces shorter contigs sequences with higher remapping rates but a significant diminution of the number of potential chimeras was observed. We conclude that the read assignation performed before the assembly step largely contributes to limit the production of chimeras. This shows that the use of SRC_c impacts the *de novo* assembled transcriptome quality and contributes to address one of the most delicate *de novo* assembly challenge [43]. The MEGAN6 contigs assignation from M2 shows more contigs than SRC_c could assign to host and symbiont (Tab. 3 and Additional files 2). In contrast, MEGAN6 assigned less contigs to host and symbiont than SRC_c for the M3. We suggest that SRC_c performs well in non-model organisms context with libraries containing taxonomically close organisms reference sequences.

### SRC_c helps us to make new biological assumptions

For all models, the SRC_c strategy led to a higher number of annotated contigs, however as only partial information on the annotation content were provided separately for the host or the symbionts in previous publications [24, 30], we were mainly restricted to qualitative comparisons. Comparing the M1a host transcriptomes to the previous study transcriptome, very few similarities were found for the most occurring functions, even if the most annotated function is common (i.e. metabolic process GO). Our 20 most occurring functions include signal transduction functions (14% of the total annotations) and molecule transport functions (8% of the total annotations) that do not appear in the most occurring function from [24]. These newly highlighted functions could help better understanding the *Orbicella faveolata* host with respect to communication and cellular exchanges with its partners. We were not able to perform a similar analysis for the symbiont transcriptome since authors of previous studies focused on the host transcriptome. For M2, only 1/15 and 3/15 common annotations were found for host and symbiont respectively. We suggest that the divergences in the analytical pipeline used, here Trinity versus CLC for *de novo* assembly followed by InterProScan versus FastAnnotator for functional annotation, make the functional annotations contents hardly comparable between studies. Despite these discrepancies, results from both analyses must be considered as potentially valuable and have to be checked with genome alignment when available or through *in vitro* validation when considering restricted group of functions (*e.g*. PCR).

Symbioses involving single cell heterotrophic hosts and photosynthetic symbionts have been described in the oceanic plankton using morphological and molecular data [5–7, 15]. Radiolarians and their symbiotic microalgae (*e.g*. Haptophytes, Dinoflagellates) have an ecological and biogeochemical significance [44–47], but little is known about symbiosis establishment and maintenance. If most microalgal symbionts can be grown in the laboratory as free-living stage [Meng et al. *submitted]*, the study of radiolarian host only relies on single-cell isolation from the field [36, 48]. In this study, the radiolarian host belongs to the Collodaria order which is ubiquitous and abundant in the open ocean [36, 49]. Our knowledge about their ecology and evolution is limited and hence our analyses represent an opportunity to learn more about the genetic repertoire of such uncultivable, non-model lineage. Regarding functional annotations, the SRC and the *noSRC* strategies provided very similar results but the SRC strategy categorized the GO annotations among 4 subsets (host, symbiont, shared and unassigned) (Additional files 5), which can be explored independently, allowing group specific interpretations and biological hypothesis building for each partner from the holobiont. For instance, symbiont CDs linked to the photosystem I and II were detected, confirming that SRC_c succeeded to assign reads to photosynthetic actors, as expected here for the symbiotic partner (Additional files 4).

### Strategies regarding the use of SRC_c and future perspectives

SRC_c successfully compared different holobiont read sets to large reference libraries in less than 24h, with reasonable computational resources (*i.e*. 10 CPUs and less than 20Go of RAM). By setting parameters (*i.e*. solidity threshold, k-mer size, similarity threshold), we adapted SRC_c to heterogeneous nature of sequences in libraries (*i.e*. length, row reads or assembled genes/transcripts, data volume, k-mers distribution) and to poorly studied systems. When studying meta-transcriptome reads, selecting abundant k-mers helps to remove the one corresponding potentially to sequencing errors; however rare sequence k-mers are consequently lost. On the contrary, when indexing already assembled sequences from genomes or transcriptomes, we do not expect a redundancy of the k-mers such as in high-throughput sequencing experiments, and we thus assume that any k-mer is relevant when it comes from a reference sequence. Accordingly, in this study, we kept the default k-mer solidity threshold value that was appropriate when indexing reads (*i.e*. sequences shorter than 300 bp, with a relatively high coverage), and lowered it to 1 when indexing longer sequences as ESTs or assembled genes. Due to the presence of small reads (50 bp) in our holobiont datasets, we also modified the default k-mer size value of 31 to a value of 25, so that any read contains at least a few k-mers. Usually the k-mer size is higher [50], however 25 base pairs corresponds to a decent value to ensure the uniqueness of the read [51]. During the query phase of SRC_c, a query sequence (from a dataset Q) must contain at least s% positions covered by at least one indexed k-mers (from a dataset B), to be considered similar to data from the set B [23]. As the s default value is set to 50%, it means that a read of size l should have at least l×s positions covered by (overlapping or nonoverlapping) indexed k-mers. Consequently, when a large majority of the reads could not be assigned, our strategy was to decrease the s parameter from 50 to 40 in order to increase the quantity of recalled reads.SRC_c implements a heuristic computing a k-mer based similarity. Contrary to BLAST-like methods, SRC_c relies uniquely on shared k-mers for its similarity computation. It means that a certain amount of error-free k-mers (*i.e*. k-mers that do not contain sequencing errors) must be found in common in order to output sequences, which can make SRC_c less sensitive compared to alignment methods which authorize mismatches. However contrary to alignment methods, SRC_c was tailored to scale to very high-volume datasets and comparisons presented in [23] showed that SRC_c could handle sets of orders of magnitudes higher volumes than BLAST (Additional files 7). SRC_c’s efficiency relies on its particular probabilistic data structure. The lightweight indexing and query of k-mers is made at the price of rare false positives. In our case, false positives correspond to k-mers that are not contained in the original indexed library. Such a false positive rate is controlled and low (Additional files 7). As in this work, the k-mer size was relatively low (i.e. 25), the default value for this parameter was kept ensuring a low false positives rate. For longer k-mers (i.e. size > 31), we recommend to increase the size of the fingerprint if more precision is needed. SRC_c can also be used in a no-false positive mode that requires more memory, but that is still less costly than a hash table as demonstrated in [23].

In our tests, SRC_c helps to retrieve holobiont reads similar to host or symbiont close species. Previous tools like COMMET [50] already proposed such computation, although their data structure makes difficult the use of k-mers of small size, as computation time would be drastically impacted. SRC_c was chosen for its simple output and its adaptability to the heterogeneous nature of the libraries studied. This is simply made by adapting the k-mer lowest occurrence and size parameters.

Future works on SRC_c parameters settings could include more extensive exploration of the impact of the similarity threshold parameter on the sensitivity of our approach. In this regard, if the reads similarity rate to the libraries could be relaxed, it may decrease the number of unassigned reads in particular for poorly studied models. A second strategy would be to implement an iterative enriching strategy to maximize the proportion of holobiont reads assigned to the host or to the symbiont. This strategy can allow to assign more sequences in the case of non-model organisms. After a first assignment round with SRC_c, holobiont reads linked to an identified group (host/symbiont) can be added to the reference libraries. Then, based on these new enriched libraries, a second run of SRC_c can be performed on the holobiont reads. This can be implemented as an iterative pipeline: at each round, more reads will be assigned to the host or symbiont categories and will then be used as reference libraries. Finally, the approach proposed here has been applied to holobiont systems (between 2 partners) but it could be used to address larger metatranscriptomic datasets composed of more complex assemblages. Depending on the SRC_c library content, the user can choose to target either one or more specific species among the variety that composed such metatranscriptomic datasets. Coupled to our assembly and downstream analysis strategy, the subsets resulting of the used of SRC_c are processed *de novo* allowing the potential discovery of newly assembled transcripts and the exploration of the functional their functional feature contents without reference genome.

## Conclusions

SRC_c successfully processed a variety of large-scale datasets and offered a pragmatic way to classify sequences from different holobiont partners before assembly. We showed that our strategy allows improving assembly metrics, and also helped to reduce drastically the proportion of chimeras in the newly *de novo* assembled sequences. Our approach offers an efficient, large scale, comparison strategy to assemble and study holobionts involving non-model organisms. Overall, this *de novo* approach, allowing a taxonomic categorization of functionalities, can reveal the link between identity and function, which is necessary to better understand the functioning and contribution of each partner in holobiont systems.

## Methods

### Radiolaria-Dinophyta holobiont model (M3) sampling, RNA-seq library and sequencing

The Collodaria colony was sampled in the South Pacific Ocean at the station 112.01 (coordinates in decimal degrees: latitude −23.3, longitude −133.9) during the *Tara* Oceans expedition in 2011 [52]. The radiolarian colony of few centimeters diameter was collected *in situ* at the subsurface (1m deep) with a plastic jar, preventing disruption of the colony and aggregation of other planktonic organisms. Live observations through the binocular were performed to verify that no organisms were accidentally attached to the colony before preservation. The collected colony was directly isolated in 15 mL of RNAlater (ThermoFisher Scientific, Waltham, MA) and preserved at −20°C. Total RNA extraction was performed using NucleoSpin RNA kit (Macherey-Nagel, Düren, Germany) starting from a slice (about 1 cm diameter) of Collodaria PAC 37 colony. Briefly, frozen cells were transferred in a 1.5 mL tube containing 100 μL RA1 lysis buffer and grinded for 1 min with a motor driven pellet pestle previously refrigerated in liquid nitrogen. Then 250 μL RA1 lysis buffer, previously mixed with 3,5 μL β-mercaptoethanol (1% of total RA1 volume), were added to the lysed cells and the total volume was transferred to a Nucleospin filter. After centrifugation and addition of an equal volume of 70% ethanol, the RNA was purified following the manufacturer’s instructions and finally eluted in 40 μL nuclease-free water. Quantity and quality of extracted RNA were assessed by capillary electrophoresis on an Agilent Bioanalyzer (Agilent Technologies, Santa Clara, CA).

Finally, in order to reduce as far as possible the risk of residual genomic DNA, a further DNase treatment was applied on the total RNA using Turbo DNA-free kit (Thermo Fisher Scientific), according to the manufacturer’s protocol. After purification with the RNA Clean and Concentrator-5 kit (ZymoResearch, Irvine, CA), RNA was eluted in 10 μL nuclease-free water and used to synthetize cDNA with the Ovation RNA-seq System Version 2 (NuGEN, San Carlos, CA), following the manufacturer’s protocol. After cDNA shearing by Covaris E210 instrument (Covaris, Woburn, MA), Illumina library was prepared using the SPRIWorks Library Preparation System on a SPRI TE instrument (Beckmann Coulter Genomics, Danvers, MA), according to the manufacturer’s protocol without size selection. Ligation products were PCR-amplified using Illumina adapter-specific primers and Platinum Pfx DNA polymerase (ThermoFisher Scientific). After library profile analysis by Agilent 2100 Bioanalyzer and qPCR quantification (MxPro, Agilent Technologies), the library was sequenced using 101 base-length read chemistry in a paired-end flow cell on HiSeq2000 Illumina sequencer (Illumina, San Diego, CA), in order to obtain nearly 50 million paired end reads.

### Data retrieval and sequence libraries construction

For each holobiont model, sequence libraries were created based on published data from taxonomically close organisms to host and symbiont species. Detailed statistics of these reference libraries can be found in Additional files 1.

For the Cnidaria-Dinophyta holobiont model (M1), the host library includes 20 assembled transcriptomes (466,582 contigs) of cnidarian organisms [53] and 2 genome-derived ESTs (201,677 ESTs) of *Nematostella vectensis* and *Orbicella faveolata* [54]. The symbiont library is composed of 123 RNA-seq reads datasets (a total of 5,563,498,607 reads) of Dinophyta from the MMETSP project [55]. We built 3 versions of the symbiont reference library, one composed of all Dinophyta (M1 a), the second exclusively composed of *Symbiodinium* spp. (15 RNA-seq datasets, a total of 123,122,726 reads) (M1 b) and the third composed of all Dinophyta except *Symbiodinium* spp. (108 RNA-seq datasets, a total of 5,440,375,881 reads) (M1 c).

For the Porifera-Bacteria holobiont model (M2), 4 RNA-seq datasets of poriferan species were included in the host library (642,229,924 total reads): *Amphimedon queenslandica* [56] *Crella elegans* [57] and both *Haliclona amboinensis* and *Haliclona tubifera* [58]. The complete bacterial gene catalog (40,154,822 assembled gene sequences) derived from the first stations from the *Tara* Oceans expedition [39] has been downloaded to constitute the symbiont reference library (OM-RGC).

For the Radiolaria-Dinophyta holobiont model (M3), we gathered Rhizaria sequences from 4 *de novo* assembled holobionts: 7,215 presumed host transcripts were extracted among a total 15,404 *de novo* assembled transcripts [15]. Host specific sequences were extracted from holobionts assemblies removing first sequences from prokaryotic origin with a blastn (e-value 1e-3) against the OM-RGC database, and second, removing symbionts sequences with a blastx (e-value 1e-3) against Dinophyta *de novo* assembled transcriptomes [Meng et al. *submitted]*. The exhaustive Dinophyta library created for the M1a was used for the reference symbiont library.

### Comparing meta-transcriptomes (i.e. holobiont reads) to reference libraries using Short Read Counter (SRC_c)

#### >Presentation of SRC_c

Short Read Connector Counter (SRC_c) [23] relies on a very lightweight data structure called a quasi-dictionary that enables to work with voluminous sequence sets. The quasi-dictionary enables to associate a piece of information to any element from a static set composed of N distinct elements. It is composed of two parts: a minimal perfect hash function (MPHF) [59] and a fingerprint table. The MPHF allows to index very efficiently the elements of the set in memory, such that each element can be associated to any piece of information (*i.e*. k-mer coverage, location in reads,…). The fingerprint table is used to verify the membership of an element to the indexed set of elements using the MPHF. This way, stranger elements to the MPHF can be filtered out. The quasi-dictionary is a probabilistic structure with a controlled false positive rate that depends on the size of the fingerprint. SRC_c needs as input two sets of sequences (that can be identical). To compare sequences from a query set Q to those from a target set T, the set indexed in the quasi-dictionary is a set of k-mers from T. Finally, for each sequence S from Q, the number of k-mers of S shared with T provides a similarity measure of S with the set T. This implies that the similarity measure given is asymmetrical: it depends on the placement of the k-mers on the reads of Q, not of those of T. SRC_c is available at https://github.com/GATB/ short read connector, the commit 94aa6a65b5ddf61eba95108069fae29c41e51fb0 was used for this study.

#### > Application on data

In this study, SRC_c is used to assign reads from an holobiont transcriptome either to the host or to the symbionts. We divided the query of the holobiont data set Q in two parts, one that consists in the comparison of Q reads to a bank (i.e. reference library) of host sequences, and another that performs the comparison to a bank of symbiont sequences. The sets to index are composed of k-mers from the sequences. In each comparison, two sequence sets are considered. The whole holobiont set Q and the target bank set B. First, the set B, which contains reads or assembled sequences and represents sequences close to the host (resp. symbiont), is indexed. During the indexation phase, the solid set of k-mers (i.e. the set composed of any k-mer which occurrence is above a user-fixed threshold (the solidity threshold) in the data set) from T is computed using the DSK [60] method. This set is next indexed in the quasi-dictionary previously described. Then the reads from the holobiont data set (Q) are queried. For each read, the query phase reports the abundance of its indexed k-mers. In the meantime, reads are checked to have enough positions (i.e. more than a given threshold which can be parameterized) for which an indexed k-mer starts over their length. This enables to add stringency to the query: a read that shares only a few k-mers with the index is considered not enough similar to the index. Finally, each read from Q (the holobiont) which was found similar to T (the host or the symbionts) during the query are returned in a binary vector and can be extracted to a FASTA format.

#### > Parameters choice

Parameters from SRC_c must be carefully chosen. First, the solidity threshold is adapted according to the nature of the sequences in the bank data set. For libraries which sequences are reads (symbiont libraries for model 1) the default value for the solidity threshold (= 2) was kept. For longer sequences (host libraries for model 1, sequences of models 2 and 3) the threshold was adapted and set to 1 when using libraries of assembled sequences or EST (host libraries for model 1, sequences of models 2 and 3). We chose a k-mer length of 25 according to the smaller input read length. We set the similarity value *s* to 50% for models 1 and 3, and decreased it to 40% for model 2. Both query and indexation phases are parallelized in SRC_c. For this study analyses were performed on a Linux system with 40 cores, with the option -t 0 (maximal number of available threads is used) and 250 GB of memory.

### Read filtering, de novo assembly and downstream analysis

All read subsets resulting from the SRC_c step were first filtered (sequences trimming and cleaning) with the Trimmomatic program [61] (v0.36) and custom parameter SLIDINGWINDOW: 10:20. Filtered reads were assembled using the *de novo* transcriptome assembly program Trinity [41] (v2.4.0) with default parameters. The newly assembled contigs metrics were calculated with the Transrate program [62] (v1.0.3). Additional downstream analyses include protein coding domain prediction using Transdecoder [63] (v3.0.1) and functional annotation with InterProScan 5 [64] (v5.24-63), both with default parameters. The pipeline used for the steps described above is publicly available on a GitHub repository https://github.com/arnaudmeng/dntap [53, Meng et al. *submitted]*.

### Taxonomic assignment with MEGAN6

The contigs sequences were compared to the nr database (August 2017 version) with the DIAMOND software [66] (v0.28.22.84) using default parameters for BLASTx comparison and a e-value of 1e^−3^. The resulting alignments were processed with the *daa2rma* tool script provided with MEGAN6 and GeneInfo Identifier (GI) were mapped to alignments using the gi_taxid.bin file (version of May 2017). Finally, taxonomic assignment has been calculated with default parameters using the MEGAN LCA (Last Common Ancestor) algorithm and were visualized through the MEGAN6 software.

### Chimeras identification

We followed the protocol described in [67]. 50,000 randomly sampled *de novo* assembled contigs for the M3 (with the SRC strategy and without SRC strategy) were compared to the 7,215 Rhizaria presumed contigs from [15] and 3,494,295 coding domains from *de novo* assembled contigs of 54 dinoflagellates transcriptomes [Meng et al. *submitted]*. The comparison was made using the BLASTx program [68] (e-value 1e^−3^). The tools scripts *detect_chimera_from_blastx.py* from [67] was applied to resulting alignments to detect potential chimeras.

## Declarations

### Acknowledgements

We thank the RCC staff for providing the dinoflagellates cultures as well as ABIMS staff for the help on computational facilities. This work was supported by a 3-year Ph.D. grant from the “Interface pour le Vivant” (IPV) program at the University of Pierre et Marie Curie (UPMC), Paris, France. This project was supported by Région Ile-de-France and benefited from the support of the project IMPEKAB ANR-15-CE02-0011.

### Author’s contributions

LB and AM designed the analysis, and LB guided the study. IP, JD and FN performed sampling and culture steps. AA and CDS optimized the molecular protocols and performed the sequencing analysis. AM and CM performed the computational analyses, with the help of PP and EC. AM, CM, PP, SLC, FN and LB wrote the manuscript. EP and PW provided critical discussions. All authors read and approved the final manuscript.

### Data Accessibility

Link to data: http://application.sb-roscoff.fr/project/radiolaria/

### Additional Files

**Additional file 1** SRC_c library content information and data sources. Table with detailed information of SRC_c libraries contents. The type of data and the total library sizes are displayed. It includes taxonomic contents and links to data repositories for holobiont models M1, M2 and M3 and data that constitute SRC_c reads/sequences libraries.

**Additional file 2** Taxonomic assignment of SRC assembled contigs with MEGAN6 for the holobiont models M1 and M2.

**Additional file 3** details of common GO annotations M1 and M2 our contigs versus previous studies

**Additional file 4** Comparison of functional annotations between SRC assembled transcriptomes and a *de novo* assembled transcriptome without the use of SRC_c in the case of holobiont model M3. Details of the functional annotations results for the SRC strategy applied to M3, the tables displayed correspond to the top 15 GO annotations found in host, symbiont, shared and unassigned transcriptomes for the three levels of annotations (MF: Molecular Functions, BP: Biological Process and CC: Cellular Component).

**Additional file 5** Radiolaria-Dinophyta meta-transcriptome taxonomic assignment with MEGAN6. Table of taxonomic assignation of the 167,023 *de novo* assembled contigs from the assembly without SRC reads sorting of the holobiont model M3.

